# The doublecortin-family kinase ZYG-8^DCLK1^ regulates motor activity to achieve proper force balance in *C. elegans* acentrosomal spindles

**DOI:** 10.1101/2023.11.22.568242

**Authors:** Emily R. Czajkowski, Nikita S. Divekar, Sarah M. Wignall

## Abstract

Although centrosomes help organize spindles in most cell types, oocytes of most species lack these structures. During acentrosomal spindle assembly in *C. elegans* oocytes, microtubule minus ends are sorted outwards away from the chromosomes where they form poles, but then these outward forces must be balanced to form a stable bipolar structure. How proper force balance is achieved in these spindles is not known. Here, we have gained insight into this question through studies of ZYG-8, a conserved doublecortin-family kinase; the mammalian homolog of this microtubule-associated protein is upregulated in many cancers and has been implicated in cell division, but the mechanisms by which it functions are poorly understood. Interestingly, we found that ZYG-8 depletion from oocytes resulted in spindles that were over-elongated, suggesting that there was excess outward force following ZYG-8 removal. Experiments with monopolar spindles confirmed this hypothesis and revealed a role for ZYG-8 in regulating the force-generating motor BMK-1/kinesin-5. Importantly, further investigation revealed that kinase activity is required for the function of ZYG-8 in both meiosis and mitosis. Altogether, our results support a model in which ZYG-8 regulates motor-driven forces within the oocyte spindle, thus identifying a new function for a doublecortin-family protein in cell division.

## INTRODUCTION

In most cell types, centrosomes function to nucleate and organize a microtubule-based spindle (Page and Hawley, 2003). However, oocytes of most species lack centrosomes and yet are able to form a bipolar spindle (Dumont and Desai, 2012; Mullen et al., 2019). In human oocytes, these acentrosomal spindles have been shown to be highly unstable; after bipolarity is achieved, the poles often split apart, and this instability leads to a high incidence of chromosome segregation errors (Holubcova et al., 2015). The mechanisms by which oocyte spindles are formed and stabilized are thus important to understand.

In recent years, *C. elegans* has emerged as a powerful model to dissect mechanisms that promote acentrosomal spindle assembly and maintenance, since rapid depletion methods have been developed that enable the removal of proteins from oocytes within minutes (Zhang et al., 2015; Divekar et al., 2021). These methods have made it possible to either deplete proteins prior to spindle formation, to probe for roles in spindle assembly, or to remove proteins from pre-formed spindles and then assess effects on the stability of the structure (Divekar et al., 2021; Cavin-Meza et al., 2022a; Cavin-Meza et al., 2022b).

In *C. elegans* oocytes, assembly of the spindle begins with nuclear envelope breakdown. Microtubules initially nucleate and form a cage-like structure around the meiotic chromosomes. Subsequently, microtubule minus ends are sorted outwards to form multiple poles that eventually coalesce into two distinct poles. Thus, to generate a stable spindle structure, forces must be produced during spindle assembly to sort microtubules outwards, and then these forces must be balanced after the spindle forms to maintain proper spindle size. KLP-18/kinesin-12, a plus-end directed microtubule motor, sorts microtubule minus ends outwards to establish bipolarity; depletion of KLP-18 results in a monopolar spindle in which microtubules minus ends coalesce into a single pole at the center with plus ends radiating outwards (Wignall and Villeneuve, 2009; Wolff et al., 2016b). KLP-18 is also required to maintain spindle bipolarity; if KLP-18 is inactivated after the bipolar spindle has already formed, the two poles converge together to create a monopolar spindle (Wolff et al., 2022b).

A recent study demonstrated that in addition to KLP-18, another plus end directed motor, BMK-1/kinesin-5, also provides outward force on the spindle. Although BMK-1 depletion does not cause defects on its own since KLP-18 provides the primary outward force (Bishop et al., 2005; Wignall and Villeneuve, 2009), a role for BMK-1 was revealed in experimental conditions where KLP-18 was depleted (Cavin-Meza et al., 2022a). Interestingly, this study also demonstrated that dynein provides an inward force on the spindle; acute dynein depletion caused the spindle to elongate and the poles to split (Cavin-Meza et al., 2022a). Thus, a balance of motor-generated outward and inward forces is essential to form and maintain the acentrosomal spindle in *C. elegans* oocytes. However, how these forces are regulated is not known.

Here, we provide insight into the mechanisms guiding force balance on the acentrosomal spindle through analysis of ZYG-8, a microtubule-associated protein that belongs to the doublecortin family of proteins. ZYG-8 contains a microtubule-binding doublecortin domain and a kinase domain (Gonczy et al., 2001). Previous studies using a series of temperature-sensitive *zyg-8* mutants demonstrated that ZYG-8 is required for proper spindle positioning in *C. elegans* mitotically dividing embryos (Gonczy et al., 2001; Bellanger et al., 2007) and for anaphase B spindle elongation in oocytes (McNally et al., 2016). However, whether ZYG-8 contributed to the assembly of the oocyte spindle was not examined. Here, we show that ZYG-8 disruption results in excess outward force on the acentrosomal meiotic spindle, and we go on to demonstrate that this protein acts to dampen the activity of motors that generate outward forces. Further, we reveal that ZYG-8’s kinase activity is required for its functions in both mitosis and meiosis, and we provide evidence that ZYG-8 plays additional roles in stabilizing spindles, beyond regulating forces.

Together, this work reveals new functions for ZYG-8 during oocyte meiosis and provides new insights into how forces are properly balanced within acentrosomal spindles.

## RESULTS

### Temperature sensitive zyg-8 mutants have oocyte spindle defects

To begin investigating the function of *C. elegans* ZYG-8 in acentrosomal spindle assembly, we utilized two different *zyg-8* temperature sensitive mutant strains, one with a mutation within the microtubule-binding Doublecortin domain, *zyg-8(or484),* and the other with a mutation within the kinase domain, *zyg-8(b235)* (**Figure 1A**, **S1A**). These mutants have been previously characterized and both display mitotic defects at the restrictive temperature of 25°C (Bellanger et al., 2007; Gonczy et al., 2001). Following incubation of these strains at this temperature, we found that oocyte spindles displayed various morphological defects (**Figure 1B-E**, **Figure S1B-E**). First, staining with the microtubule minus end marker ASPM-1 revealed pole defects; while spindles at the permissive temperature had two ASPM-1 clusters, an increased number of spindles at the restrictive temperature had 3 or more clusters, reflecting that the spindles were multipolar and/or had fragmented poles (18/48 *or484* spindles and 24/62 *b235* spindles; **Figure 1E**, **S1E**). Spindles at 25°C were also longer than control spindles (**Figure 1C**, **S1C**), and many spindles appeared to be bent. To quantify this phenotype, we used ASPM-1 staining to define the center point of each pole. We then drew lines connecting each pole to the middle of the spindle, defined by the center of the DNA staining, and measured the angle between the two poles (**Figure 1B** schematic); a perfectly straight spindle would therefore measure 180 degrees. Using this measurement, we found that spindles at the restrictive temperature were on average significantly more bent than at the permissive temperature (**Figure 1D**, **S1D**). Together, these results suggest that ZYG-8 is required for proper spindle assembly in oocytes.

**Figure 1.**
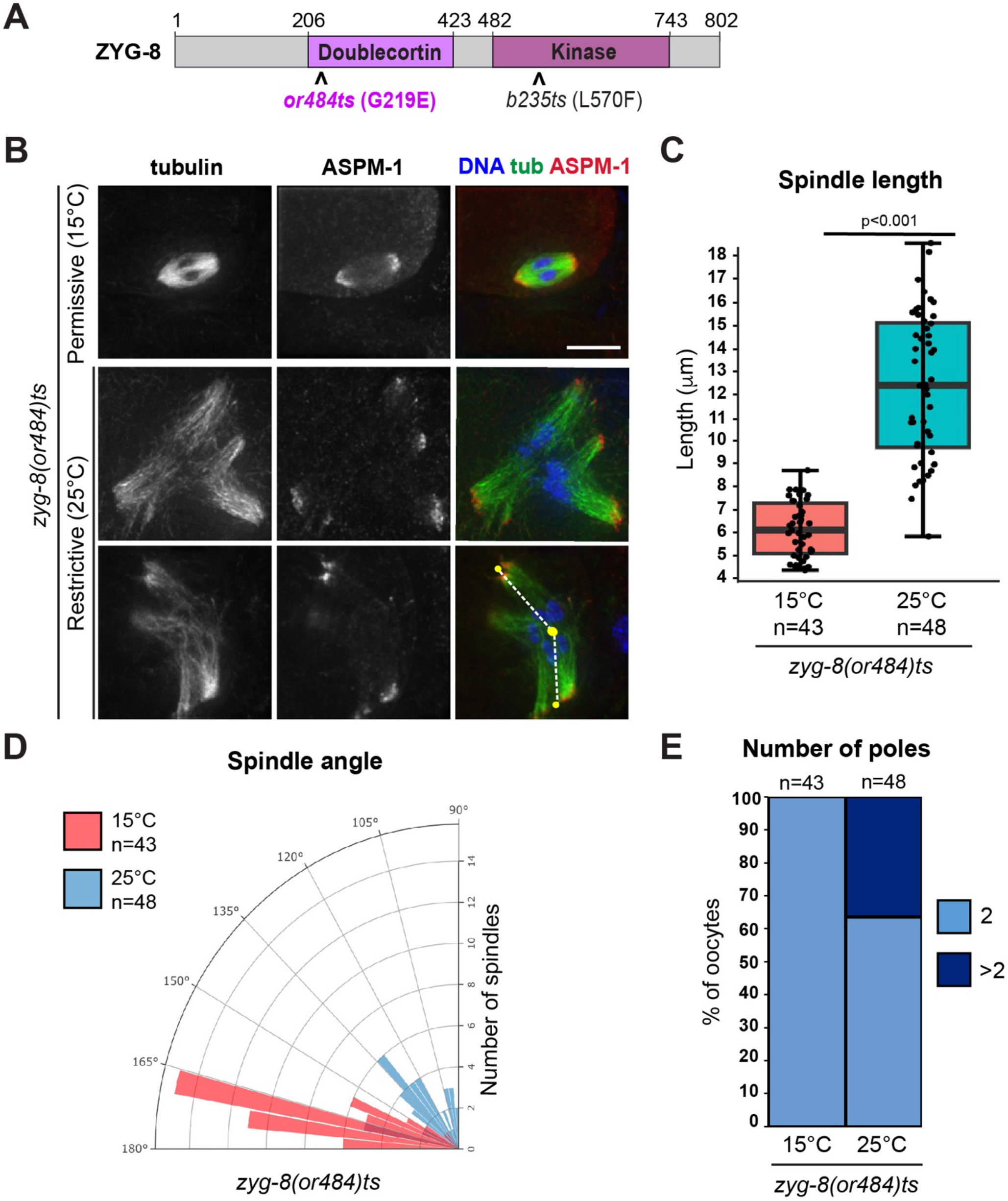
A *zyg-8* temperature sensitive mutant displays defects in oocyte spindle morphology at the restrictive temperature. (A) ZYG-8 schematic, highlighting the microtubule binding doublecortin domain, the kinase domain, and the location of the *or484* temperature sensitive mutation. (B) Immunofluorescence images of *zyg-8(or484)* oocytes at either the permissive (15°C) or restrictive (25°C) temperatures. Shown are tubulin (green), DNA (blue), and ASPM-1 (red). Lines drawn on the bottom right image represent how spindle angles were measured (for the quantification shown in panel D). (C-E) Quantification of spindle length, spindle angle, and number of ASPM-1-marked poles. After incubation at the restrictive temperature, oocyte spindles were on average longer, more bent, and some had additional ASPM-1-marked poles. Scale bar = 5µm.

### ZYG-8 localizes diffusely across the meiotic spindle and is required for proper spindle assembly

To investigate ZYG-8 further, we turned to the auxin-inducible degradation (AID) system, which enables spatially and temporally controlled depletion of proteins (Zhang et al., 2015). Using CRISPR/Cas9 technology, we introduced a GFP::degron tag onto the N-terminus of ZYG-8 in a strain expressing the ubiquitin ligase TIR1 from a germline-specific promoter (hereafter referred to as “ZYG-8 AID”; **Figure 2A**). To validate this strain, we incubated adult worms on auxin containing plates for 18 hours (“long-term AID”) and verified that this treatment resulted in substantial ZYG-8 depletion from the germ line using embryo-only western blotting (**Figure S2A**). Notably, under these conditions spindles in mitotically-dividing one-cell stage embryos were mispositioned (15/15 spindles), phenocopying defects observed in previous studies of *zyg-8* mutants (**Figure S2B**) (Gonczy et al., 2001; Bellanger et al., 2007). Moreover, we analyzed the ZYG-8 AID strain in the absence of auxin, to determine if tagging ZYG-8 caused major phenotypes on its own. Notably, we did not detect any embryonic lethality in the ZYG-8 AID strain in the absence of auxin (<1% dead eggs) and the lengths and angles of oocyte spindles were similar to a control strain that expresses TIR1 but has untagged ZYG-8 (**Figure S2C-D**). Therefore, we set out to use this strain to study the role of ZYG-8 in oocytes.

**Figure 2.**
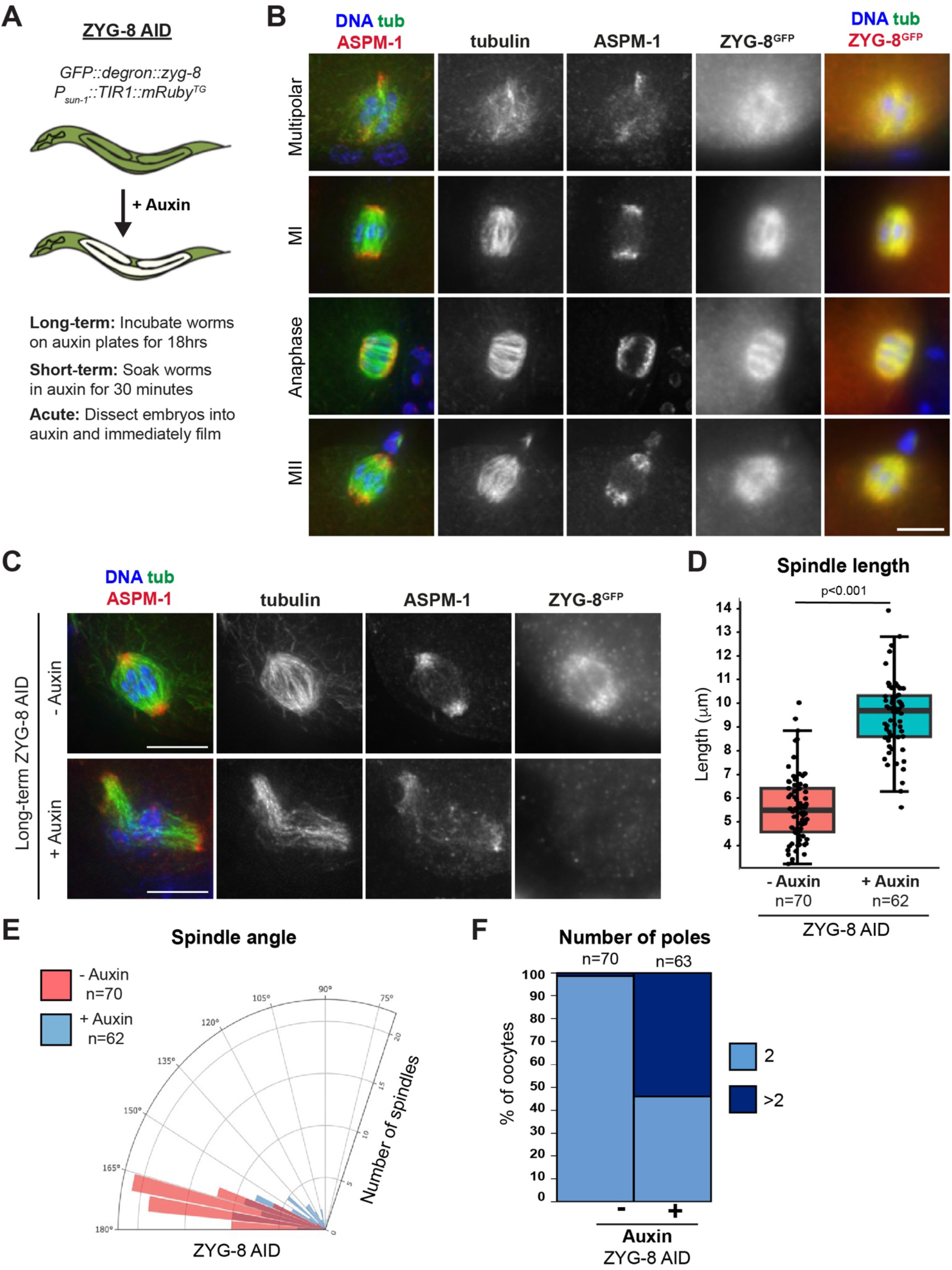
ZYG-8 localizes diffusely across the oocyte spindle and is required for proper spindle assembly. (A) Schematic representation of the auxin-inducible degradation system for ZYG-8 depletion in *C. elegans.* (B) Immunofluorescence images of oocytes in the ZYG-8 AID strain showing ZYG-8 localization throughout meiosis; shown are tubulin (green), DNA (blue), ASPM-1 (red, left panel), and ZYG-8 (stained using a GFP antibody; red, right panel). ZYG-8 is diffusely localized to the oocyte spindle throughout Meiosis I and II. Due to diffuse ZYG-8 localization, ZYG-8 images shown are not deconvolved. (C) Immunofluorescence images of either control spindles or spindles treated overnight on auxin-containing plates (“long-term AID”); shown are tubulin (green), DNA (blue), ASPM-1 (red), and ZYG-8 (stained using a GFP antibody). (D-F) Quantification of spindle length, spindle angle, and number of ASPM-1-marked poles, demonstrating that long-term ZYG-8 depletion causes spindle defects. Scale bars = 5µm.

Because ZYG-8 is tagged with GFP in the ZYG-8 AID strain, we first assessed ZYG-8 localization throughout all stages of meiosis in oocytes. ZYG-8 localization to microtubules begins during spindle assembly, with ZYG-8 localized diffusely across the entire meiotic spindle. ZYG-8 remains on the spindle through anaphase, and this pattern repeats in Meiosis II (**Figure 2B**).

Notably, ZYG-8 staining was undetectable upon long-term AID, again confirming robust depletion (**Figure 2C**). Under these conditions, spindles were longer than they were in the absence of auxin (**Figure 2D**). Moreover, auxin-treated spindles were on average more bent than control spindles (**Figure 2E**) and some spindles contained more than two ASPM-1 clusters (**Figure 2F**), mirroring the phenotypes seen in the temperature-sensitive *zyg-8* mutants (**Figure 1**, **S1**). These results supported the hypothesis that ZYG-8 is required for normal spindle formation in oocytes and also confirmed that the ZYG-8 AID strain could be used to further probe ZYG-8 function.

### ZYG-8 is required to maintain the stability of pre-formed oocyte spindles

The spindle elongation and bending observed in response to ZYG-8 disruption could be consistent with excess outward force on the spindle. During spindle assembly, forces are generated to push microtubule minus ends outwards and then once bipolarity is achieved, these forces must be balanced to maintain proper spindle size. Therefore, an increase in outward forces following ZYG-8 depletion could cause the observed increase in spindle length and might also cause the poles to split and the center of the spindle to lose integrity and bend.

To investigate this hypothesis, we sought to remove ZYG-8 from pre-formed spindles to test whether there was evidence for activation of outward forces. To this end, we generated a version of the ZYG-8 AID strain expressing GFP::tubulin and mCherry::histone; using this strain we could dissect oocytes directly into auxin and visualize the effects of ZYG-8 depletion on the spindle in real time (“acute AID”). To enrich for pre-assembled bipolar spindles, we arrested oocytes in Metaphase I by depleting anaphase promoting complex (APC) component EMB-30. Without auxin, metaphase-arrested spindles in ZYG-8 AID oocytes maintain bipolarity (**Figure 3A, Video 1,** 5/5 videos). In contrast, upon dissection into auxin, ZYG-8 AID oocyte spindles immediately began to elongate and lose mid-spindle integrity (**Figure 3A, Video 2**; phenotypes seen in 10/10 videos), consistent with an activation of outward forces.

**Figure 3.**
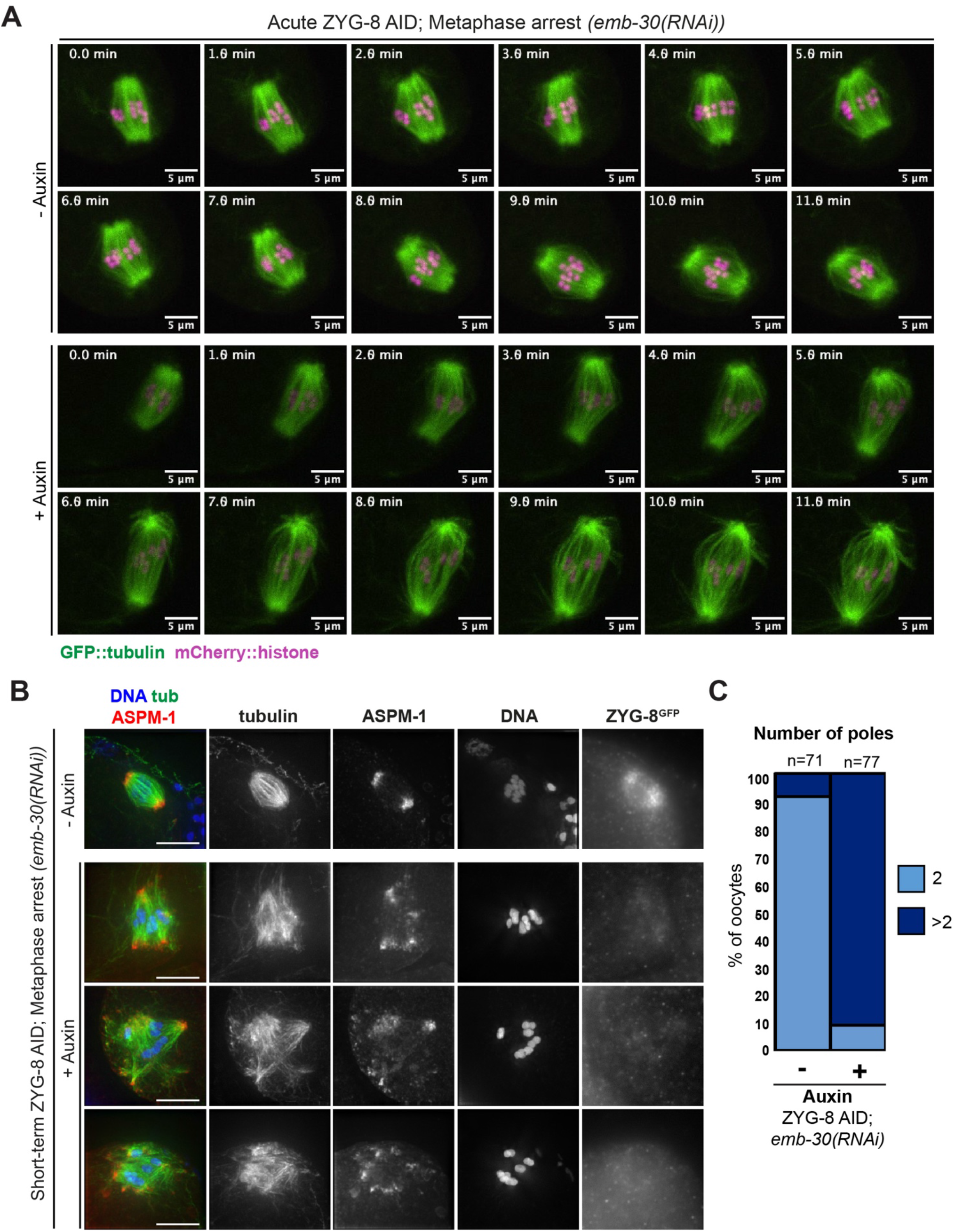
ZYG-8 depletion from pre-formed spindles results in spindle elongation and loss of bipolarity. (A) Live imaging of *emb-30(RNAi)* metaphase-arrested spindles; shown are GFP::tubulin (green) and mCherry::histone (magenta). In the absence of auxin (top panels), spindles are able to maintain bipolarity and chromosome alignment throughout the time-lapse. In auxin treated oocytes (“acute AID”, bottom panels), the spindle elongates, the midspindle weakens, and chromosomes become misaligned. (B) Immunofluorescence images of metaphase-arrested *(emb-30(RNAi))* spindles in the ZYG-8 AID strain. When worms were soaked in auxin for 30-40 minutes (“short-term AID”; rows 2-4), spindles became multipolar and chromosomes appeared more dispersed compared to control spindles (row 1). (C) Quantification of number of poles per spindle in control and short-term AID metaphase-arrested spindles. Scale bars = 5µm.

To assess the consequences of more extended auxin treatment on pre-formed spindles, we soaked *emb-30(RNAi)* ZYG-8 AID worms in auxin for 30 minutes and then performed immunofluorescence (“short-term AID”). Under these conditions, we observed severe spindle defects **(Figure 3B**). Following auxin treatment, over 80% of spindles had 3 or more ASPM-1 clusters, reflecting pole fragmentation and/or loss of spindle bipolarity (**Figure 3C**); the severity of this phenotype precluded the measurement of spindle lengths, since it was difficult to define a primary spindle axis in most cases. Additionally, in contrast to control metaphase arrested oocytes, we noticed that chromosomes were often misaligned after short-term ZYG-8 AID **(Figure 3B**).

To ensure that these spindle phenotypes were not artifacts caused by the *emb-30(RNAi)*-induced metaphase arrest, we performed both short-term and acute ZYG-8 depletion on unarrested oocytes. Following short-term AID depletion, we observed an increased number of spindles with more than two ASPM-1 clusters and we observed a greater average spindle angle and length compared to untreated worms (**Figure S3A-D**). Moreover, we also observed defects when we performed acute ZYG-8 AID on unarrested oocytes. While control spindles were able to maintain bipolarity and undergo anaphase, oocytes treated with auxin exhibited defects, including destabilization of the midspindle and spindle bending (**Figure S3E, Video 3, 4**). Altogether, these data demonstrate that ZYG-8 is required for both the assembly and stability of acentrosomal spindles. Moreover, the observed phenotypes are consistent with activation of outward forces.

### ZYG-8 depletion triggers excess outward force in the absence of KLP-18

Our hypothesis that outward forces are activated upon ZYG-8 depletion raised the possibility that this protein may normally suppress a factor that generates these forces. Since KLP-18/kinesin-12 provides the major outward sorting force in *C. elegans* oocytes (Wolff et al., 2016; Wolff et al., 2022b), we first sought to explore the possibility that ZYG-8 regulates KLP-18. To this end, we co-depleted KLP-18 and ZYG-8, to determine if there was evidence for excess outward force under these conditions.

Following *klp-18(RNAi)*, microtubule minus ends are not sorted outwards during spindle assembly, so a monopolar spindle forms with microtubule minus ends in the center and microtubule plus ends oriented towards the periphery of the aster (Wignall and Villeneuve, 2009; Wolff et al., 2016) (**Figure 4A**, **4C**). When we combined *klp-18(RNAi)* with long-term ZYG-8 AID, monopolar spindles were still able to form, with microtubule minus end marker ASPM-1 concentrated at the single pole. However, we noticed that ASPM-1 was often also present at the periphery of the spindle on the ends of microtubule bundles, suggesting that some microtubule minus ends were sorted outwards during spindle assembly (**Figure 4A**, **4C**). We observed the same phenotype when we performed *klp-18(RNAi)* on the *zyg-8(or484)* temperature-sensitive mutant and then incubated it at the restrictive temperature; monopolar spindles formed that contained ASPM-1-marked minus ends at the periphery (**Figure 4B**, **4C**). Thus, ZYG-8 inactivation enables some microtubule minus ends to be sorted outwards during spindle assembly in oocytes lacking KLP-18, suggesting that some outward forces were being generated in this condition.

**Figure 4.**
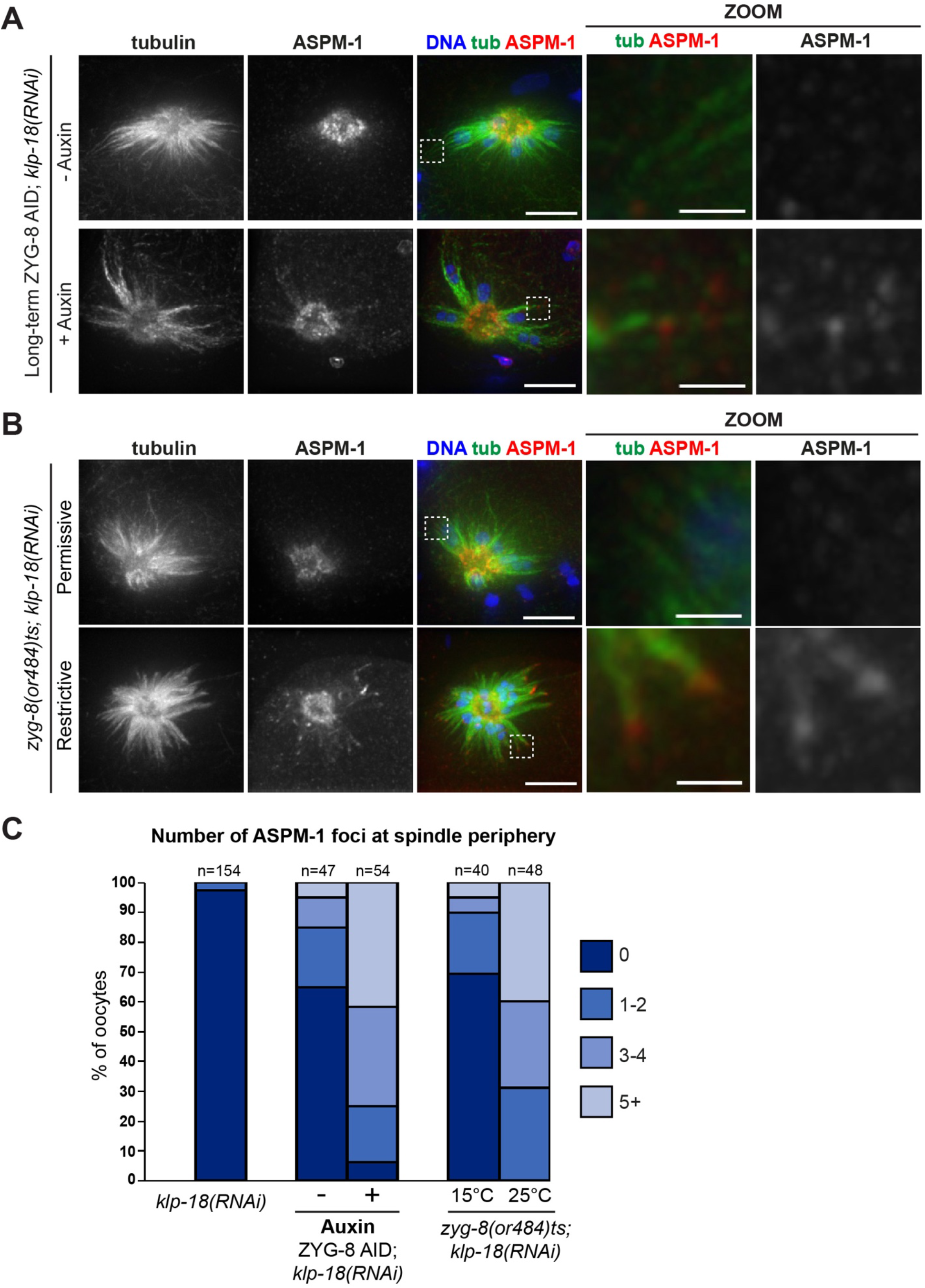
ZYG-8 depletion causes excess outward sorting in the absence of KLP-18. (A) Immunofluorescence images of *klp-18(RNAi)* oocytes in the ZYG-8 AID strain. Shown are tubulin (green), DNA (blue), and ASPM-1 (red). In the absence of auxin, a monopolar spindle forms with ASPM-1-marked microtubule minus ends concentrated in the center (top row). Upon long-term AID to deplete ZYG-8, a central ASPM-1-marked pole still forms but there is also ASPM-1 localization at the end of microtubule bundles at the spindle periphery (zoom), suggesting that minus ends are being sorted outwards despite the loss of KLP-18. (B) Immunofluorescence images of *klp18(RNAi)* oocytes at either the permissive (15°C) or restrictive (25°C) temperatures in the *zyg-8(or484)* temperature sensitive mutant. Shown are tubulin (green), DNA (blue), and ASPM-1 (red). After incubation at the restrictive temperature, monopolar spindles have ASPM-1 localized to the periphery of the spindle at the end of microtubule bundles (zooms). Scale bar = 5µm (regular images); 1µm (zooms). (C) Quantification of number of ASPM-1 foci seen at the periphery of each monopolar spindle.

Next, we sought to remove ZYG-8 from pre-formed *klp-18(RNAi)* monopolar spindles to determine whether this would result in the activation of outward forces. Remarkably, when we performed short-term ZYG-8 AID on *klp-18(RNAi)* monopolar spindles, we saw a loss of monopole integrity in most oocytes (78% of total) and spindles were sometimes able to restore bipolarity (40% of total; **Figure 5A**, **5B**). We observed the same phenotype using live imaging. In the absence of auxin, *klp-18(RNAi)* spindles maintain a single monopole as chromosomes move towards the center of the aster during anaphase (Muscat et al., 2015) (**Figure 5C, Video 5**). In contrast, following acute ZYG-8 AID monopolar spindles began to reorganize (12/12 videos), and these spindles were sometimes able to reestablish bipolarity and segregate chromosomes bidirectionally (4/12 videos; **Figure 5C, Video 6, 7).** Thus, ZYG-8 depletion results in the activation of outward forces in the absence of KLP-18, suggesting that ZYG-8 may regulate another force-generating factor.

**Figure 5.**
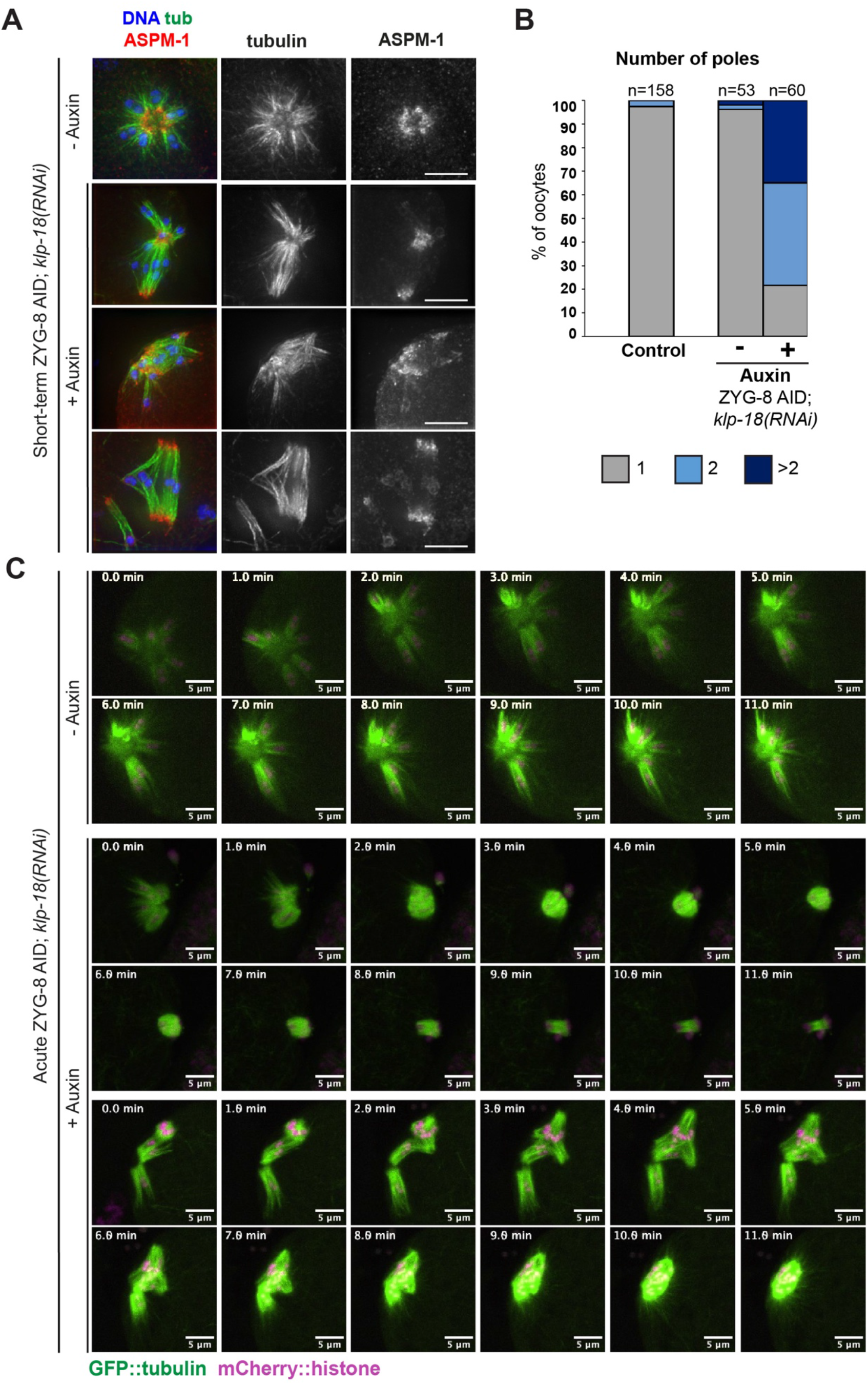
Removal of ZYG-8 from pre-formed monopolar spindles triggers spindle reorganization and can restore bipolarity. (A) Immunofluorescence images of ZYG-8 AID; *klp-18(RNAi)* oocytes. Shown are tubulin (green), DNA (blue), and ASPM-1 (red). Upon short-term auxin treatment to deplete ZYG-8 from preformed monopolar spindles, microtubule minus ends were no longer organized into a single pole, suggesting that microtubule minus ends were sorted outwards; in some cases, bipolarity appeared to be largely restored (bottom row). (B) Quantification of spindle polarity from the experiment shown in A. (C) Live imaging of *klp-18(RNAi)* ZYG-8 AID spindles; shown are GFP::tubulin (green) and mCherry::histone (magenta). Control monopolar spindles remain monopolar as chromosomes slowly move towards the center pole in anaphase (rows 1-2). When treated with auxin, monopolar spindles reorganize (rows 3-6). In some cases, these spindles were able to segregate chromosomes bidirectionally, suggesting that bipolarity was restored (rows 3-4). We also observed cases where a monopolar MII spindle reorganized and appeared to incorporate the polar body, ultimately forming what appeared to be a bipolar spindle (rows 5-6). Scale bars= 5µm.

### With loss of BMK-1 and KLP-18, outward forces are no longer activated upon ZYG-8 depletion

Although KLP-18 is the major outward force generating motor in *C. elegans* oocytes, we recently reported that BMK-1/kinesin-5 provides a redundant outward sorting force that is not detected when KLP-18 is present (Cavin-Meza et al., 2022a). Additionally, BMK-1 is broadly localized across the meiotic spindle through MI and MII (Bishop et al., 2005; Cavin-Meza et al., 2022a), similar to ZYG-8 (**Figure 2B**). Therefore, we reasoned that ZYG-8 may be functioning to inhibit BMK-1. To investigate this hypothesis, we generated a version of the ZYG-8 AID strain that also contained a loss of function *bmk-1(ok391)* mutation (Bishop et al., 2005). We then performed *klp-18(RNAi)* combined with short-term ZYG-8 AID, to determine if outward forces were still activated in this assay if BMK-1 function was compromised. Strikingly, with the loss BMK-1, monopoles remained intact following ZYG-8 depletion (**Figure 6A**, **6B**), suggesting that outward forces were not activated. This phenotype was also confirmed using live imaging, where auxin-treated ZYG-8 AID monopolar spindles did not reorganize into bipolar spindles in the *bmk- 1(ok391)* mutant and we never observed chromosomes segregating bidirectionally in anaphase; instead chromosomes moved towards the center of the monopole, indistinguishable from normal monopolar anaphase (**Figure 6C, Video 8, 9, 11;** 5/5 auxin-treated spindles remained monopolar). Together, these data suggest that ZYG-8 functions to inhibit BMK-1.

**Figure 6.**
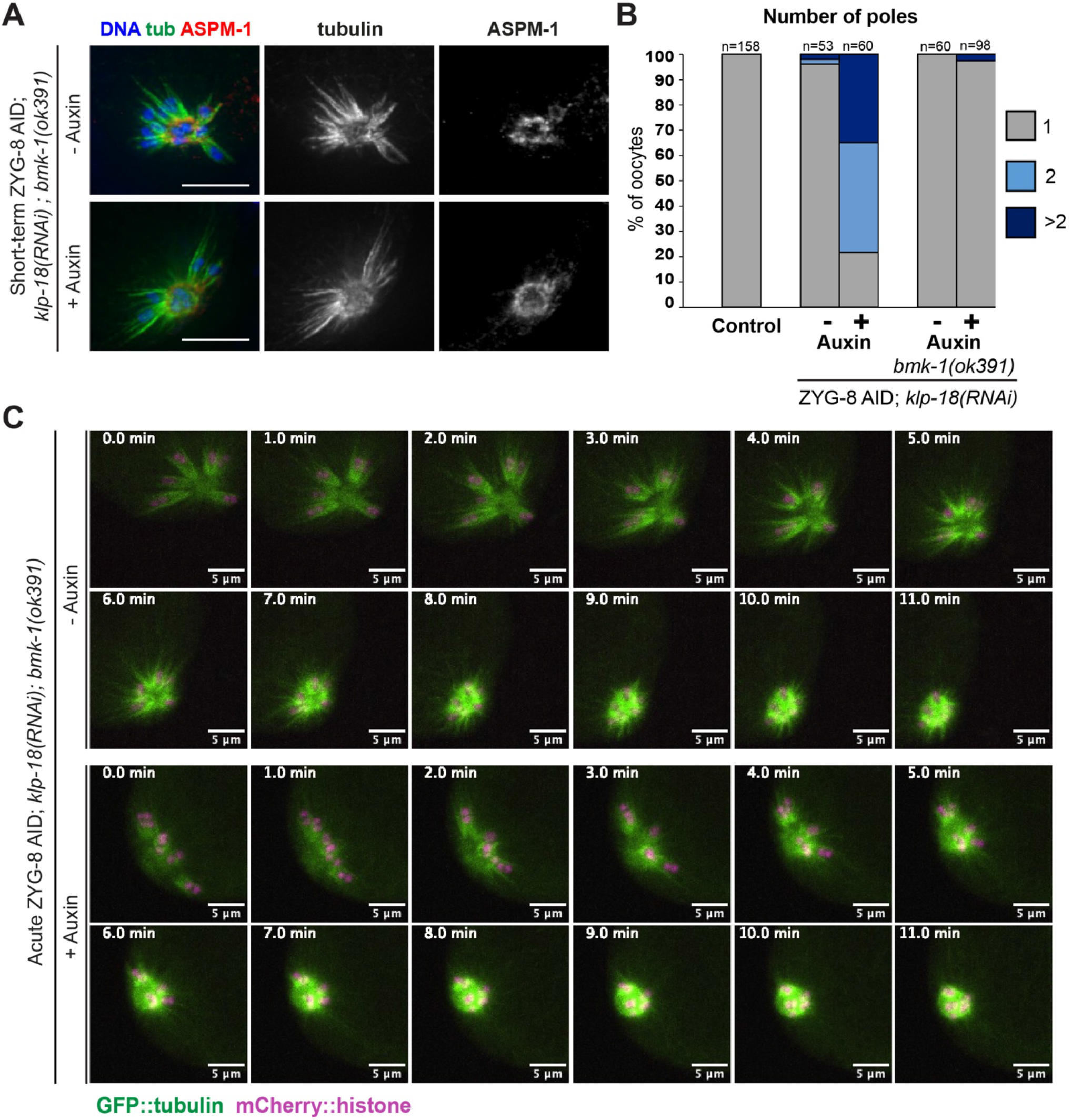
With loss of BMK-1, monopolar spindles do not reorganize upon ZYG-8 removal. (A) Immunofluorescence images of *klp-18(RNAi)* in ZYG-8 AID; *bmk-1(ok391)* oocytes. Shown are tubulin (green), DNA (blue), and ASPM-1 (red). In this strain, monopolar spindles did not reorganize upon short-term auxin treatment. (B) Quantification of the experiment shown in A. Note that the data from the ZYG-8 AID strain without the *bmk-1* mutation is repeated from Figure 5B, for purposes of comparison. (C) Live imaging of *klp-18(RNAi)* in the ZYG-8 AID; *bmk-1(ok391)* strain; shown are GFP::tubulin (green) and mCherry::histone (magenta). In both the presence and absence of auxin, spindles remain monopolar as chromosomes slowly move towards the center pole in anaphase and the monopolar spindle shrinks. Scale bars = 5µm.

Although the complete rescue of monopole integrity following triple depletion of KLP-18, ZYG-8, and BMK-1 provided evidence that ZYG-8 regulates BMK-1, it did not rule out the possibility that ZYG-8 may normally regulate KLP-18 or other factors as well. To test this, we assessed the effects of depleting ZYG-8 in the *bmk-1(ok391)* strain without depleting KLP-18. Interestingly, we found that spindles were longer in the ZYG-8 AID; *bmk-1(ok391)* strain in the presence of auxin, suggesting that outward forces became overactive upon ZYG-8 depletion (**Figure S4A**, **S4B**). Moreover, some spindles had fragmented and/or multiple poles (**Figure S4C**). These results demonstrate that ZYG-8 does not solely function to regulate BMK-1. Thus, ZYG-8 likely regulates multiple factors to produce proper force balance within the spindle.

### ZYG-8’s kinase activity is required for its function in meiosis and mitosis

Finally, we sought to understand more about the mechanisms by which ZYG-8 regulates spindle forces. The mammalian homolog of ZYG-8, DCLK1 (doublecortin-like kinase 1), is upregulated in multiple cancers and there are ongoing efforts to generate inhibitors targeting this kinase as a therapeutic strategy (Bai et al., 2003; Burgess and Reiner 2000; Francis et al., 1999; Gleeson et al., 1998; Jean et al., 2012). However, whether the kinase activity of DCLK1 or any of its paralogs is required during cell division is not known. Therefore, to determine if ZYG-8’s kinase activity is required for its function, we created a ZYG-8 kinase-dead mutant strain using CRISPR. We mutated a conserved aspartic acid within the catalytic loop of ZYG-8 to asparagine; this mutation has been previously shown to render DCLK1 kinase dead (D511N in DCLK1; D604N in ZYG-8) (**Figure 7A**) (Chen and Cole, 2015; Patel et al., 2016; Patel et al., 2021). Homozygous mutant worms carrying this mutation laid 98% dead eggs, demonstrating that this mutation causes embryonic lethality.

**Figure 7.**
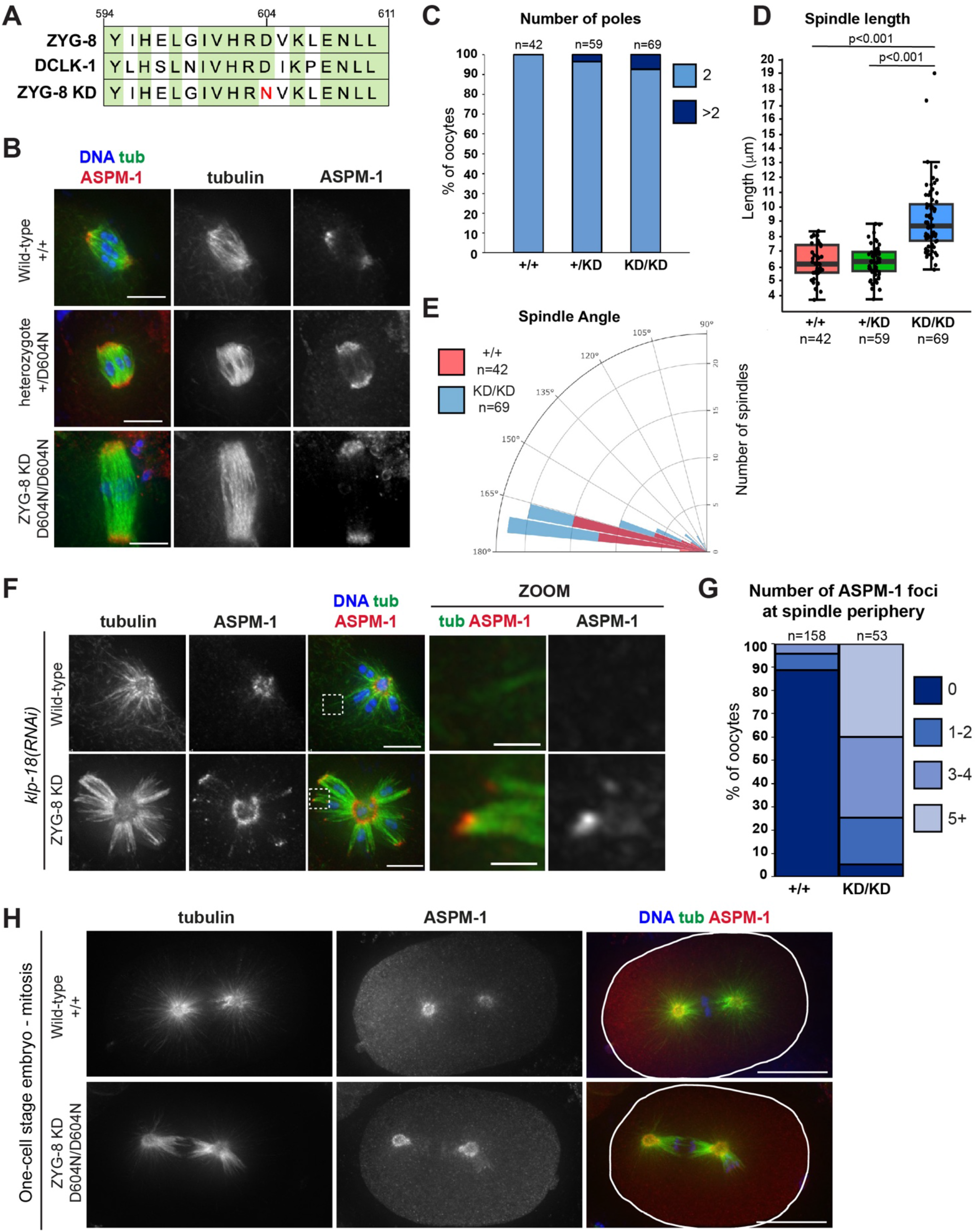
Kinase activity is required for ZYG-8 function in meiosis and mitosis. (A) Sequence of a portion of the ZYG-8 kinase domain, indicating the D604N kinase dead mutation (ZYG-8 KD). Green represents homology between ZYG-8 and its mammalian homolog DCLK1. (B) Immunofluorescence images of spindles in wild-type oocytes (row 1) and oocytes from *zyg-8^D604N^* heterozygous (row 2) and homozygous (ZYG-8 KD, row 3) parents; shown are tubulin (green), DNA (blue), and ASPM-1 (red). Oocyte spindles from wild-type and *zyg-8^D604N^* heterozygote parents were indistinguishable while spindles from *zyg-8^D604N^*homozygous parents were longer in length. (C-E) Quantification of the number of ASPM-1-marked poles, spindle length, and spindle angle in the experiment shown in B. (F) Immunofluorescence images *of klp-18(RNAi)* monopolar spindles in oocytes from *zyg-8^D604N^* homozygote parents (ZYG-8 KD, row 2) compared to wild type (row 1); shown are tubulin (green), DNA (blue), and ASPM-1 (red). ASPM-1 was present at the end of microtubules bundles at the periphery of monopolar spindles in the ZYG-8 KD strain (zooms). (G) Quantification of the number of ASPM-1 foci at the monopolar spindle periphery. (H) Immunofluorescence images of one-cell stage mitotic spindles in embryos from wild-type worms (row 1) and from *zyg-8^D604N^* homozygous parents (row 2). Spindles in the ZYG-8 KD mutant are mis-positioned, phenocopying defects seen in *zyg-8* mutants (Gönczy et al., 2001; Bellanger et al., 2007) and following ZYG-8 AID. Scale bars in B and F = 5µm (regular images); 1µm (zooms). Scale bar in H = 10µm.

To assess the effect of this mutation on meiotic spindle formation, we compared spindle morphology in oocytes from *zyg-8^D604N^* heterozygote and homozygote parents to wild-type worms. While oocyte spindles in heterozygotes were approximately the same length as wild-type spindles, homozygous *zyg-8^D604N^* oocyte spindles were substantially longer (**Figure 7B, 7D**). Similarly, when we performed *klp-18(RNAi)*, monopolar spindles were able to form in homozygous *zyg-8^D604N^*oocytes, but over 90% of the spindles had ASPM-1 staining at the periphery of the aster (**Figure 7F, 7G**). These phenotypes were reminiscent of the spindle assembly phenotypes we observed in the *zyg-8* temperature-sensitive mutant strains and following long-term ZYG-AID (**Figure 1C**, **S1C**, **2D**, **4**), suggesting that ZYG-8’s kinase activity is required to inhibit outward forces in acentrosomal oocyte spindles. Notably, we also observed mitotic defects in one-cell stage embryos from *zyg-8^D604N^*homozygous parents (**Figure 7H**). Bipolar spindles formed but they were mispositioned (10/10 *zyg-8^D604N^* spindles), phenocopying defects seen in previous analysis of *zyg-8* temperature sensitive mutants (Gonczy et al., 2001; Bellanger et al., 2007) and following ZYG-8 AID (**Figure S2B**). Thus, kinase activity is required for ZYG-8 function in mitosis as well.

Interestingly, however, we noticed that spindles in homozygous *zyg-8^D604N^* oocytes were not as severely disrupted as those in the *zyg-8ts* or ZYG-8 AID strains. We did not observe substantial fragmentation of spindle poles in the *zyg-8^D604N^*mutant (**Figure 7C**) and the spindle angles were similar to wild type (**Figure 7E**). This suggests that while kinase activity is required to dampen outward forces and prevent spindle over-elongation, other regions of the protein may function to stabilize the bipolar spindle structure to prevent pole and mid-spindle destabilization. Thus, we propose that ZYG-8 plays multiple important roles in regulating proper spindle architecture.

## DISCUSSION

### ZYG-8 is required for proper force balance in the acentrosomal spindle

Altogether, our data reveals an important role for ZYG-8 in facilitating the assembly and stability of the oocyte spindle and supports a model in which ZYG-8 functions to regulate motor-driven forces (**Figure 8**). During spindle assembly, microtubule minus ends must be pushed outwards away from the chromosomes, necessitating the production of outward forces (Wolff et al., 2016). However, mechanisms must also exist to limit these outward forces, so that the spindle can remain stable after it forms (**Figure 8A**). Previous studies revealed that KLP-18/kinesin-12 and BMK-1/kinesin-5 produce outward forces, while dynein produces a counter-balancing inward force  Wolff et al., 2016; Cavin-Meza et al., 2022a; Wolff et al., 2022b). We have now identified ZYG-8 as another factor that is important for this force balance.

**Figure 8.**
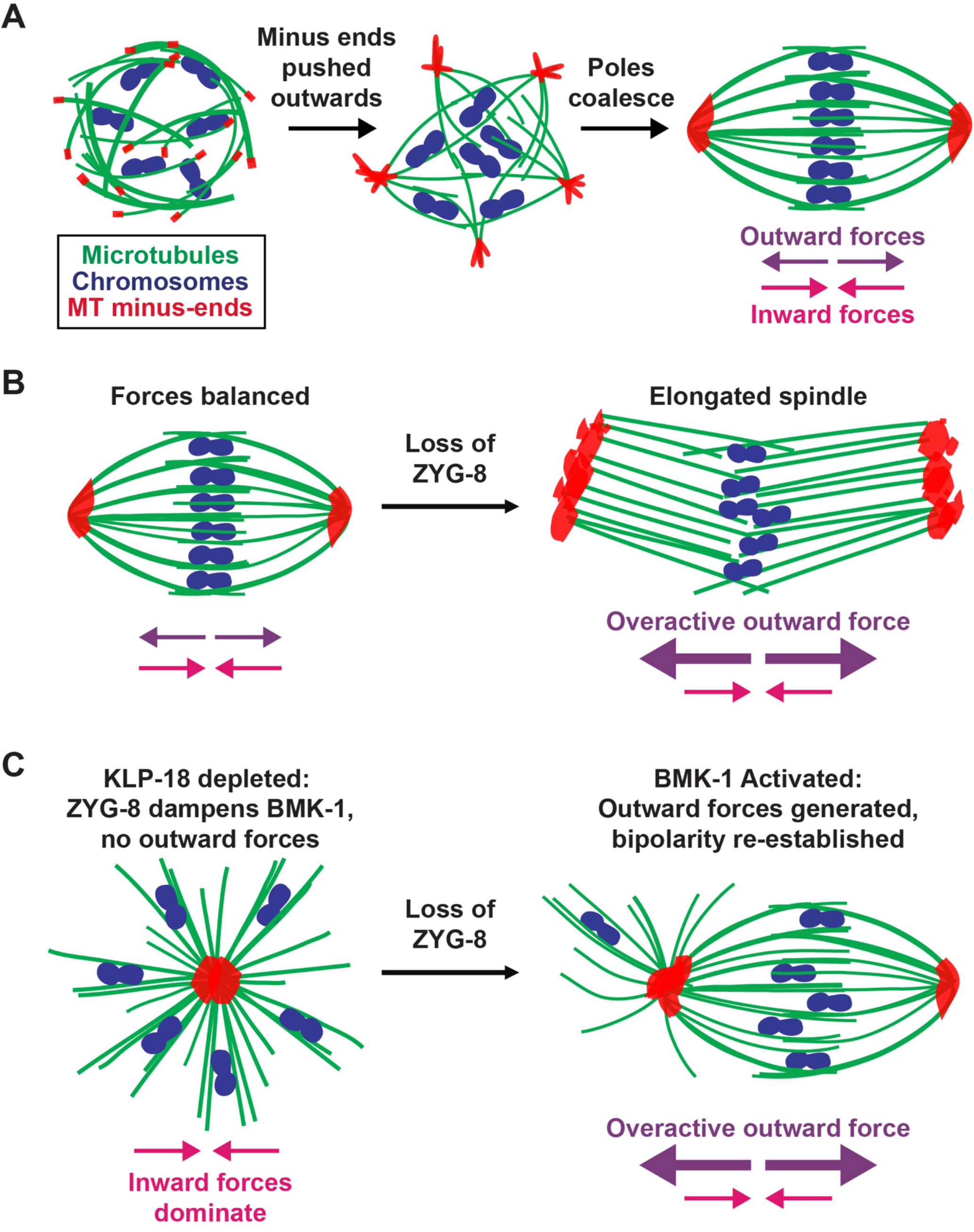
Model: ZYG-8 is required for proper force balance in the acentrosomal spindle. Chromosomes (blue), microtubules (green), and microtubule minus ends (red). (A) As an acentrosomal spindle forms, microtubules begin to nucleate in the presence of condensed chromosomes. KLP-18 (kinesin-12) provides the major outward force and sorts microtubule minus ends to the outer periphery of the forming spindle. Minus ends then form multiple poles that coalesce into two distinct poles at either end of the spindle. In a bipolar spindle, the outward and inward forces are balanced. (B) Upon removal of ZYG-8 from a balanced metaphase spindle, outward forces become overactive, pushing minus ends outward, and elongating the spindle. (C) When ZYG-8 is removed from a monopolar spindle lacking KLP-18, BMK-1 becomes over-activated and microtubule minus ends are sorted to the periphery of the spindle, reestablishing bipolarity.

A fundamental role for ZYG-8 in regulating force balance is supported by multiple findings. First, spindles formed in the absence of ZYG-8 are longer than wild type, and acute removal of ZYG-8 from pre-formed spindles causes the spindle to elongate, consistent with overactivation of outward forces (**Figure 8B**). While theoretically these phenotypes could also be caused by the dampening of inward forces or by the lengthening of spindle microtubules, experiments with monopolar spindles do not support those interpretations. Specifically, we found that ZYG-8 removal from pre-formed monopoles resulted in minus ends being pushed outwards to form multiple poles (**Figure 8C**); this dramatic rearrangement would not be expected to happen if inward forces were simply dampened or if microtubules grew longer. Remarkably, in this experiment, monopolar spindles were sometimes able to convert back to bipolar spindles that could mediate bi-directional chromosome segregation, demonstrating that the outward forces activated upon ZYG-8 depletion were sufficient to drive bipolarity. Further, we obtained evidence that ZYG-8 regulates at least one outward force-generating motor, BMK-1. When we removed ZYG-8 from monopolar spindles in *bmk-1* mutants, the monopoles did not reorganize. Thus, BMK-1 provided the outward force that drove spindle bipolarization upon ZYG-8 depletion, suggesting that ZYG-8 normally dampens BMK-1. However, we also found that ZYG-8 does not solely function to regulate BMK-1, since spindle length still increased when ZYG-8 was depleted in a *bmk-1* mutant without KLP-18 depletion. Thus, we propose that ZYG-8 regulates multiple motor-driven forces in the oocyte spindle.

In addition to spindle elongation, we also observed pole defects upon ZYG-8 depletion; spindles formed in the absence of ZYG-8 sometimes had fragmented or multiple poles. These defects are similar to those previously observed upon dynein depletion, which caused spindle poles to become broad and unfocused (Crowder et al., 2015; Cavin-Meza et al., 2022a). This raises the possibility that ZYG-8, like dynein, may play a direct role in stabilizing acentrosomal poles. However, we do not favor this interpretation for a few reasons. First, while dynein removal does not disrupt the central microtubule overlap region of the spindle (Cavin-Meza et al., 2022a), ZYG-8 depletion causes mid-spindle bending, suggesting that ZYG-8 does not solely function at poles. Second, monopolar spindles were able to form following double depletion of KLP-18 and ZYG-8; these spindles had an organized central pole, suggesting that ZYG-8 is not required for pole formation *per se*. Finally, when ZYG-8 was removed from pre-formed monopolar spindles, the microtubule minus ends that were pushed outwards were able to form poles. This phenotype stands in stark contrast to that seen following dynein depletion; when dynein was removed from pre-formed monopolar spindles, the entire monopole came apart, dispersing meiotic chromosomes throughout the cytoplasm (Cavin-Meza et al., 2022a). Thus, we infer that ZYG-8 is unlikely to play the same role as dynein in stabilizing poles directly, and instead we suggest that it plays a more global role in forming and stabilizing acentrosomal spindles.

Interestingly, the phenotypes we observed were more severe when we removed ZYG-8 from pre-formed spindles than when we depleted ZYG-8 prior to spindle assembly. Spindles formed in the absence of ZYG-8 often had split poles or were multipolar, consistent with a role for ZYG-8 in spindle assembly. However, the percentage of multipolar spindles was even higher when ZYG-8 was removed from metaphase-arrested spindles; we speculate that the arrest may be exacerbating this phenotype by pausing spindles in metaphase where they may be subjected to prolonged spindle forces. Similarly, monopolar spindles formed in the absence of ZYG-8 had some minus ends at the periphery of the spindle, but an organized central pole was able to form. However, removal of ZYG-8 after monopole formation caused spindle reorganization, both with and without a metaphase arrest. This suggests that ZYG-8 may play a more significant role in stabilizing the metaphase spindle than it does in forming the spindle. Since ZYG-8 and BMK-1 have a similar localization pattern during meiosis, we suggest that ZYG-8 loads onto the spindle along with BMK-1, and functions to inhibit BMK-1 from over elongating microtubules during metaphase.

Intriguingly, in *C. elegans* oocytes and mitotically-dividing embryos, BMK-1 inhibition causes faster spindle elongation during anaphase, suggesting that BMK-1 normally functions as a brake to slow spindle elongation (Saunders et al., 2007; Laband et al., 2017). Further, ZYG-8 has been shown to be required for spindle elongation during anaphase B (McNally et al., 2016). Our findings may provide an explanation for this phenotype, since if ZYG-8 inhibits BMK-1 as we propose, then following ZYG-8 depletion, BMK-1 could be hyperactive, slowing anaphase B spindle elongation.

### How might ZYG-8 regulate motor-driven forces?

An important question is how ZYG-8 functions to regulate force balance. Notably, our studies demonstrate that kinase activity is required for ZYG-8 function. This is an important finding, since the mammalian homolog of ZYG-8 (DCLK1/doublecortin-like kinase 1) is upregulated in a wide range of cancers, and knockdown studies have suggested that it is required for tumor progression. Because of this, DCLK1 has recently become a target for the development of kinase inhibitors as therapeutics (Chandrakesan et al., 2017; Chhetri et al., 2022; Ferguson et al., 2020; Ge et al., 2021; Westphalen et al., 2017; Westphalen et al., 2016; Weygant et al., 2014) . However, whether kinase activity is essential for DCLK1’s function in cell division was not previously known. Here, we show that kinase activity is important for the function of a DCLK1-family protein during both mitosis and meiosis in an *in vivo* model, providing evidence that may validate this therapeutic strategy.

How kinase activity regulates spindle function is an important area of future study. One possibility is that ZYG-8 phosphorylates motors directly. There is a large amount of evidence, spanning multiple species and cell types, that kinesin motors can be regulated by phosphorylation (Sato-Yoshitake et al., 1992; Matthies and Reymann, 1993; McIlvain et al., 1994; Hirokawa et al., 2009; Verhey and Hammond, 2009; DeBerg et al., 2013). For example, phosphorylation of Kinesin-5 motors can impact localization, motor activity, or function in mitosis (Blangy et al., 1995; Mann and Wadsworth, 2019). In *C. elegans*, Aurora B kinase (AIR-2) is required for BMK-1’s localization to mitotic and meiotic spindle microtubules *in vivo* and can phosphorylate BMK-1’s tail domain *in vitro*, suggesting that phosphorylation may regulate its localization (Bishop et al., 2005). In *Xenopus laevis* egg extract, Cdk1 phosphorylation increases kinesin-5 binding to microtubules (Cahu et al., 2008) and in *D. melanogaster,* kinesin-5 was shown to undergo a reversable phosphorylation in order to activate or deactivate the motor (Bickel et al., 2017). It has also been shown that several kinesin families can be regulated by autoinhibition controlled by phosphorylation; this may be a general mechanism for kinesin regulation (Verhey and Hammond, 2009). For example, in *Xenopus*, Cenp-E (kinesin-7) autoinhibition is reversed via phosphorylation of the tail domain (Espeut et al., 2008). Given these established roles for phosphorylation in modulating motor activity, it is possible that ZYG-8 directly phosphorylates BMK-1 and/or other motors directly to regulate force balance.

Alternatively, it is possible that ZYG-8 regulates motors indirectly through its microtubule binding activity. In mammals, DCLK1 and its paralog DCX (doublecortin) are required for neuronal development and can regulate motor activity in neurons (Gleeson et al., 1998; Francis et al., 1999; Burgess and Reiner, 2000; Bai et al., 2003; Jean et al., 2012). Neurons lacking DCX or DCLK1 show defects in kinesin-mediated vesicle transport in the absence of defects in microtubule organization (Deuel et al., 2006; Liu et al., 2012). Moreover, DCLK1 coats dendritic microtubules and is required for kinesin-3-mediated cargo transport into dendrites; this led to the hypothesis that crosstalk between DCLK1 and motors at the microtubule surface represented a regulatory mechanism for motor-driven cargo transport (Lipka et al., 2016). Thus, it is also possible that in *C. elegans* oocytes, microtubule-bound ZYG-8 may regulate the association of motors with the microtubule lattice, affecting their activity. Future studies will be important to fully understand how ZYG-8 and its paralogs regulate motor activity.

### Additional roles for ZYG-8 in spindle function

Importantly, our work also suggests that ZYG-8 may have additional roles beyond regulating force balance. We found that the *zyg-8^D604N^*kinase dead mutant had elongated bipolar spindles and had microtubule minus ends at the periphery of monopolar spindles, consistent with an activation of outward forces. However, spindles in this mutant did not display some of the other defects we observed in the *zyg-8* temperature sensitive mutants or following ZYG-8 depletion; *zyg-8^D604N^* spindles largely had organized poles and were not significantly bent. This implies that the pole and midspindle disruption observed following ZYG-8 depletion was not solely caused by an activation of outward forces, suggesting that ZYG-8 has additional roles in stabilizing the spindle structure.

One possibility is that ZYG-8 may promote microtubule stability to facilitate spindle assembly. Notably, DCLK-1 has been shown to phosphorylate a microtubule associated protein (MAP7D1) to facilitate axon elongation in mouse cortical neurons (Koizumi et al., 2017); it is possible that ZYG-8 could phosphorylate microtubule regulatory factors in *C. elegans* oocytes in a similar manner. Moreover, doublecortin-family proteins have been shown to regulate aspects of microtubule dynamics, including decreasing the catastrophe frequency and increasing the nucleation rate, producing a net stabilization of microtubules (Bechstedt and Brouhard, 2012; Horesh et al., 1999; Jean et al., 2012; Moores et al., 2004; Moores et al., 2006; Patel et al., 2016; Patel et al., 2021). Thus, it is possible that ZYG-8, via its microtubule-binding doublecortin domain, may directly promote microtubule stability. This hypothesis is consistent with our findings, as depletion of ZYG-8 would cause microtubule destabilization, leading to disruption of the midspindle and spindle poles, while the kinase-dead mutant, which should have the doublecortin domain intact, would not be expected to display these phenotypes. Thus, we suggest that in addition to regulating motor activity through its kinase domain, ZYG-8 also promotes microtubule stability within the oocyte spindle, likely via its microtubule-binding doublecortin domain. Further experiments investigating the mechanistic function of the doublecortin domain could further elucidate this hypothesis.

In summary, our studies have provided new insights into the functions of a doublecortin-family protein in cell division. Moreover, our work has provided new insights into how forces are balanced within acentrosomal spindles, and into how the bipolar spindle structure is stabilized. Further studies elucidating the relationship between all force generating proteins including KLP-18, BMK-1, Dynein, and ZYG-8 should be performed to better understand the relationship between these proteins and how they work together to create a bipolar spindle with properly balanced forces.

## MATERIALS AND METHODS

### C. elegans strains

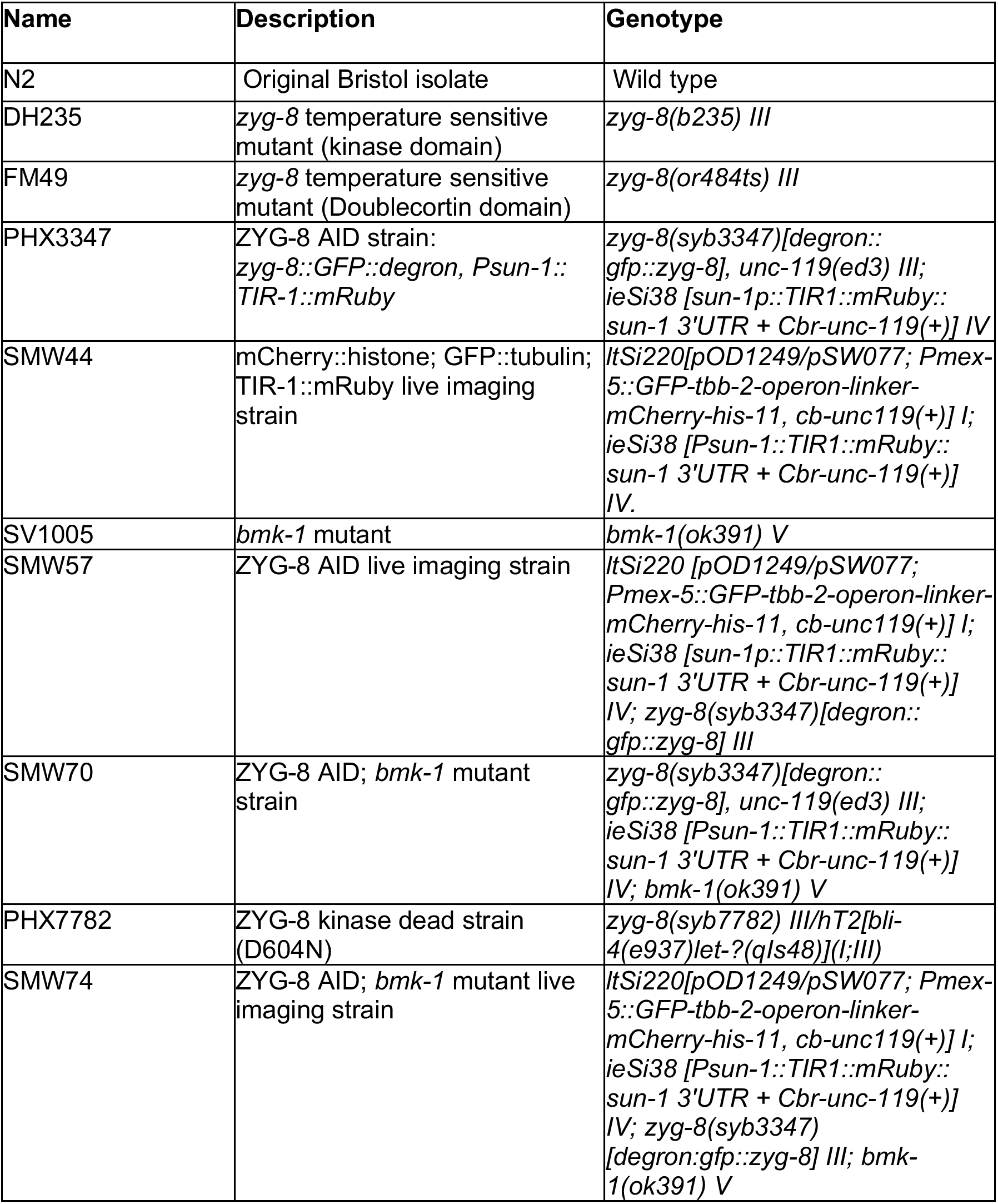

### Generation of *C. elegans* strains and maintenance

PHX3347 and PHX7782 were generated by SunyBiotech using CRISPR/Cas9 editing of the endogenous locus of *zyg-8*. All strains were maintained at 15°C. The *zyg-8^D604N^* mutation was embryonic lethal when homozygous, so the strain was balanced with the hT2 balancer. Homozygotes were over 98% embryonic lethal.

### Immunofluorescence

Immunofluorescence was performed as described in (Wolff et al., 2022a). Briefly, adult worms were picked into a 10 μL drop of Meiosis Medium (0.5 mg/mL Inulin, 25 mM HEPES, and 20% FBS in Leibovitz’s L-15 Media) (Laband et al., 2018) on a poly-L-lysine-coated glass slide. Worms were dissected to release oocytes, a coverslip was placed on top, and then the slide was plunged into liquid nitrogen and the cover slip was flicked off. Slides were fixed in -20°C methanol for 30-40 minutes (40 minutes for strains expressing GFP or mCherry in order to quench the fluorescence), washed in PBS, and then blocked in AbDil (PBS with 4% BSA, 0.1% Triton-X-100, 0.02% sodium azide) at room temperature for at least 30 minutes. Primary antibodies were diluted in AbDil and incubated at 4°C overnight. The next day, samples were washed 3X with PBS-T (PBS with 0.1% Triton-X-100) and incubated in secondary antibody diluted in AbDil for 2 hours. Samples were again washed 3X with PBS-T and incubated in mouse anti-α-Tubulin-FITC (Sigma) diluted 1:500 in Abdil at room temperature for 1 hour. Samples were rinsed again and then incubated at room temperature in Hoechst diluted 1:1000 in PBS-T for 15 minutes. Samples were then rinsed one final time in PBS-T and samples were mounted in mounting media (0.5% p-phenylenediamine, 20 mM Tris-Cl, pH 8.8, 90% glycerol) then sealed with nail polish and stored at 4°C. Slides were imaged within 2 weeks of production. Primary antibodies used were rabbit-α-ASPM-1 (1:5000, gift from Arshad Desai), rabbit-α-BMK-1 (1:500, gift from Jill Schumacher), mouse-α-Tubulin-FITC (1:500, DM1α, Sigma), and mouse-α-GFP (1:250, 3E6, Invitrogen). Alexa Fluor secondary antibodies (Invitrogen) were used at 1:500. Fixed imaging was performed on DeltaVision Core deconvolution microscope with 100X (NA = 1.4) objective (Applied Precision). This microscope is housed in the Northwestern University Biological Imaging Facility supported by the NU Office for Research. Image stacks were obtained at 0.5μm z-steps and deconvolved using SoftWoRx (Applied Precision).

### RNAi feeding

RNAi was performed as described in (Wolff et al., 2022a). Briefly, individual RNAi clones from an RNAi library (Kamath et al., 2003), were grown in LB containing 100µg/µl AMP overnight at 37°C. After 18 hours, cultures were pelleted for 10 minutes, and excess LB was removed. The pellet was resuspended in the remaining LB and spotted on agar plates containing 100µg/µl AMP and 1mM IPTG. Plates were kept in the dark at room temperature to dry overnight. Worms were synchronized by bleaching adult worms, collecting embryos, and allowing them to hatch without food overnight. These starved L1s were then plated onto the RNAi plates and grown to adulthood at 15°C for 5 days.

### Auxin treatment

#### Long-term Auxin treatment

NGM plates were prepared with auxin added to a final concentration of 1mM. Plates were stored in the dark and used within 8 weeks. 18 hours prior to fixation, L4 worms were picked onto an auxin-containing plate and left at 15°C. For experiments requiring both RNAi and auxin treatment, NGM plates were made containing 100µg/µl AMP, 1mM IPTG, and 1mM auxin, the plates were seeded with RNAi clones, and RNAi was performed as described above.

#### Short-term Auxin treatment

Meiosis Medium containing 5mM auxin was prepared using a stock of 400mM auxin dissolved in 100% EtOH. Adult worms were picked into this auxin solution and kept in a humidity chamber at room temperature for 30-45 minutes. Worms were then dissected and immunofluorescence was performed. As a vehicle control, EtOH was diluted in Meiosis Medium and the same incubation conditions were used.

### Ex Utero Live Imaging

Two-color live imaging was performed using a Nikon SoRa spinning disk confocal microscope with an oil-immersion 60x (1.42 NA) objective lens. A Yokogawa CSU-W1 dual-disk spinning disk unit with 50μm pinholes and a Hamamatsu ORCA-Fusion Digital CMOS Camera were used for image acquisition. The microscope was controlled using Nikon imaging software NIS-Elements software. The SoRa microscope is housed in the Northwestern University Biological Imaging Facility supported by the NU Office for Research.

10-15 worms were taken from desired experimental plates and dissected into 10µl of Meiosis Medium (described above) either with or without 500µM auxin. Quickly, a small Vaseline ring was made around the sample and an 18x18mm coverslip was laid on top. The slide was imaged immediately. Fifteen z-stacks at 1μm increments were taken every 20 seconds at room temperature. Images were processed using ImageJ. Images are shown as maximum intensity projections of the entire spindle structure.

### Western Blotting

For long-term auxin treatment, adult worms were transferred to either four control NGM or four NGM auxin-containing plates (100 worms/plate) then incubated for 24 hours. These plates were then washed into an Eppendorf tube (4 plates/tube) and bleached to remove worm bodies and leave only embryos. Bleached samples were spun at 800rcf for 1 minute, rinsed with M9 and spun again, and then liquid was removed, leaving the bleached embryo sample in 10µl M9. For short-term auxin treatment, 4 NGM plates of approximately 100 adult worms each were washed into an Eppendorf tube with 500µl of Meiosis Medium containing either 5mM auxin or vehicle. The worms were incubated with intermittent tube rotation for 40 minutes without light exposure. After 40 minutes, worms were bleached as described above leaving 10µl of M9 and embryo sample. 10µl samples were then mixed with 10µl SDS sample buffer, boiled at 95°C for 10 minutes, and then stored at -20°C immediately.

Samples were run on 4–20% gradient Tris-Glycine gel (BioRad Mini-PROTEAN TGX) at 80V for 2 hours. The protein was wet transferred onto nitrocellulose at 120V for 22 hours. The blot was blocked in 5% milk with TBS-0.1%Tween for 2 hours at room temperature on a rocker and then incubated with indicated primary antibodies overnight at 4°C (1:1000 mouse anti-degron (Sigma-Aldrich) or 1:5000 mouse anti-tubulin (ThermoFisher)). The blot was washed with TBS-0.1% Tween and incubated with goat anti-mouse HRP (1:5000) (ThermoFisher) for 2 hours at room temperature, washed once more, incubated for 5 minutes in BioRad Clarity ECL solution, and then exposed for 10 minutes.

### Data Analysis and quantification

#### Spindle length

To determine spindle length, three-dimensional renderings of spindles were generated using Imaris 3D Imaging Software (Bitplane). The center point of each pole was determined using the Surfaces tool to define the volume of the ASPM-1 staining, and the center point of this volume was assigned. Length was then measured as the center to center of each pole.

#### Spindle angle

The angle of the spindle was measured using Imaris Software. The poles were defined as described above. To determine the midspindle, the Surfaces tool was used to define the volume of the DNA signal and the center point of this volume was assigned; this was then used as the vertex while the poles were used as the arms, and the inside angle of the spindle was measured.

#### Number of poles

Quantifications were performed by counting poles per spindle as defined by ASPM-1 staining. A pole was defined as ASPM-1 staining at the end of MT bundles, with an accumulation of more than 3 ASPM-1 foci.

#### Number of ASPM-1 foci at spindle periphery

Quantifications were performed by counting the number of ASPM-1 foci at the end of MT bundles at the periphery of monopolar spindles.

### Statistical Methods

Statistical analysis was performed using R studio; a Shapiro-Wilks test was used to confirm data distribution normality. Normally distributed data was analyzed using a two-tailed t-test, while non-normally distributed data was analyzed using a Mann-Whitney Wilcoxon non-parametric test.

## ACKNOWLEDGMENTS

We would like to thank members of the Wignall lab and the WiLa ICB for support and guidance, with a particular thanks to Gabe Cavin-Meza, Hannah Horton, Juhi Narula, and Jordy Martinez for comments on the manuscript. We are also grateful to Arshad Desai for antibodies and the *Caenorhabditis* Genetics Center, funded by NIH Office of Research Infrastructure Programs (P40 OD010440), for strains. This work was supported by Cellular and Molecular Basis of Disease (CMBD) Training Grant NIH T32 GM00806 (to ERC) and NIH R01GM124354 and NIH R01GM141386 (to SMW). Microscopy was performed at the Biological Imaging Facility at Northwestern University, supported by the Chemistry for Life Processes Institute, the NU Office for Research, and the Department of Molecular Biosciences.

---

**Figure S1.**
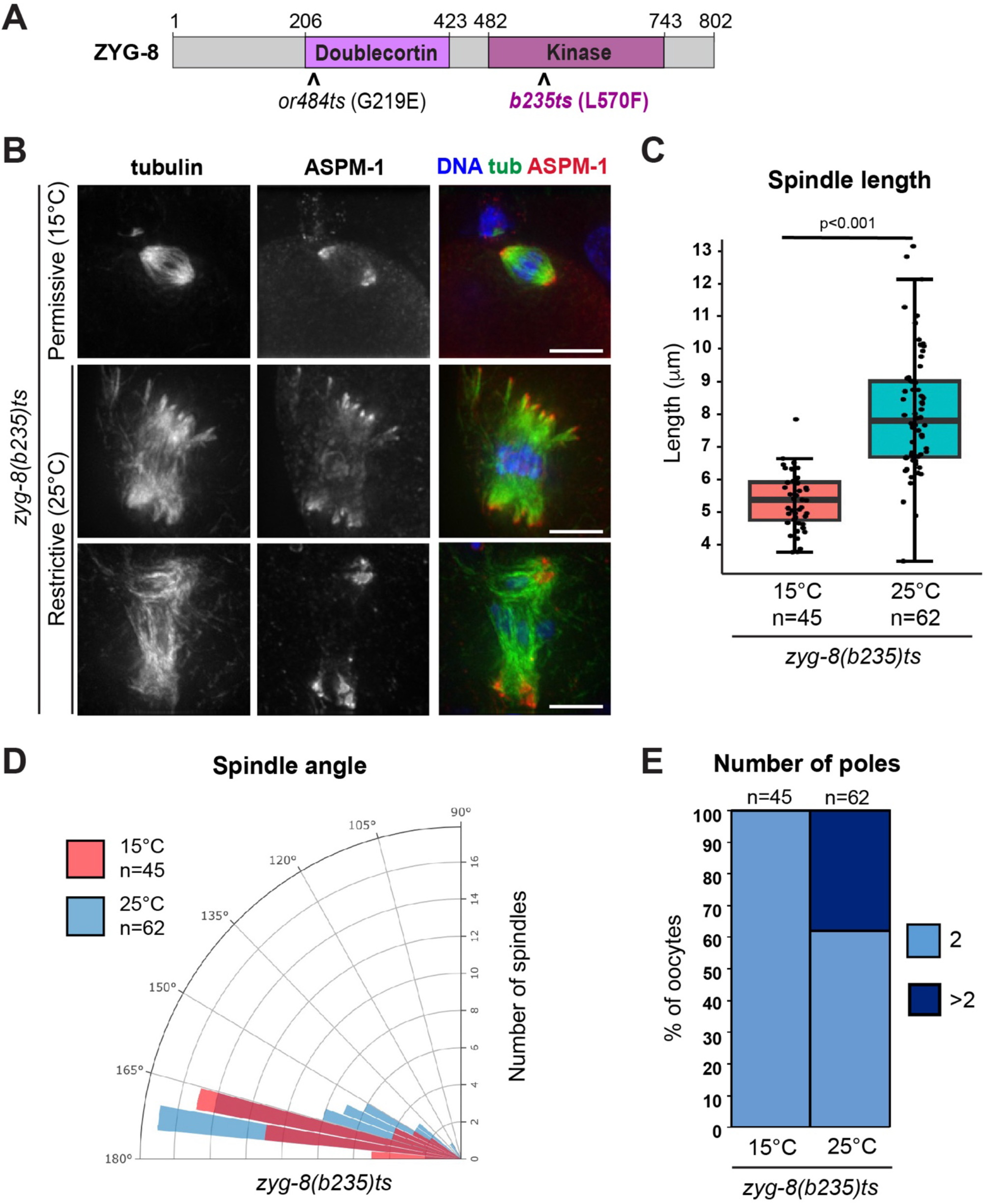
A *zyg-8* temperature sensitive mutant displays defects in oocyte spindle morphology at the restrictive temperature. (A) ZYG-8 schematic, highlighting the microtubule binding doublecortin domain, the kinase domain, and the location of the *b235* temperature sensitive mutation. (B) Immunofluorescence images of *zyg-8(b235)* oocytes at either the permissive (15°C) or restrictive (25°C) temperatures. Shown are tubulin (green), DNA (blue), and ASPM-1 (red). (C-E) Quantification of spindle length, spindle angle, and number of ASPM-1-marked poles in the experiment shown in B. After incubation at the restrictive temperature, oocyte spindles were on average longer, more bent, and some had additional ASPM-1-marked poles. Scale bars = 5µm.

**Figure S2.**
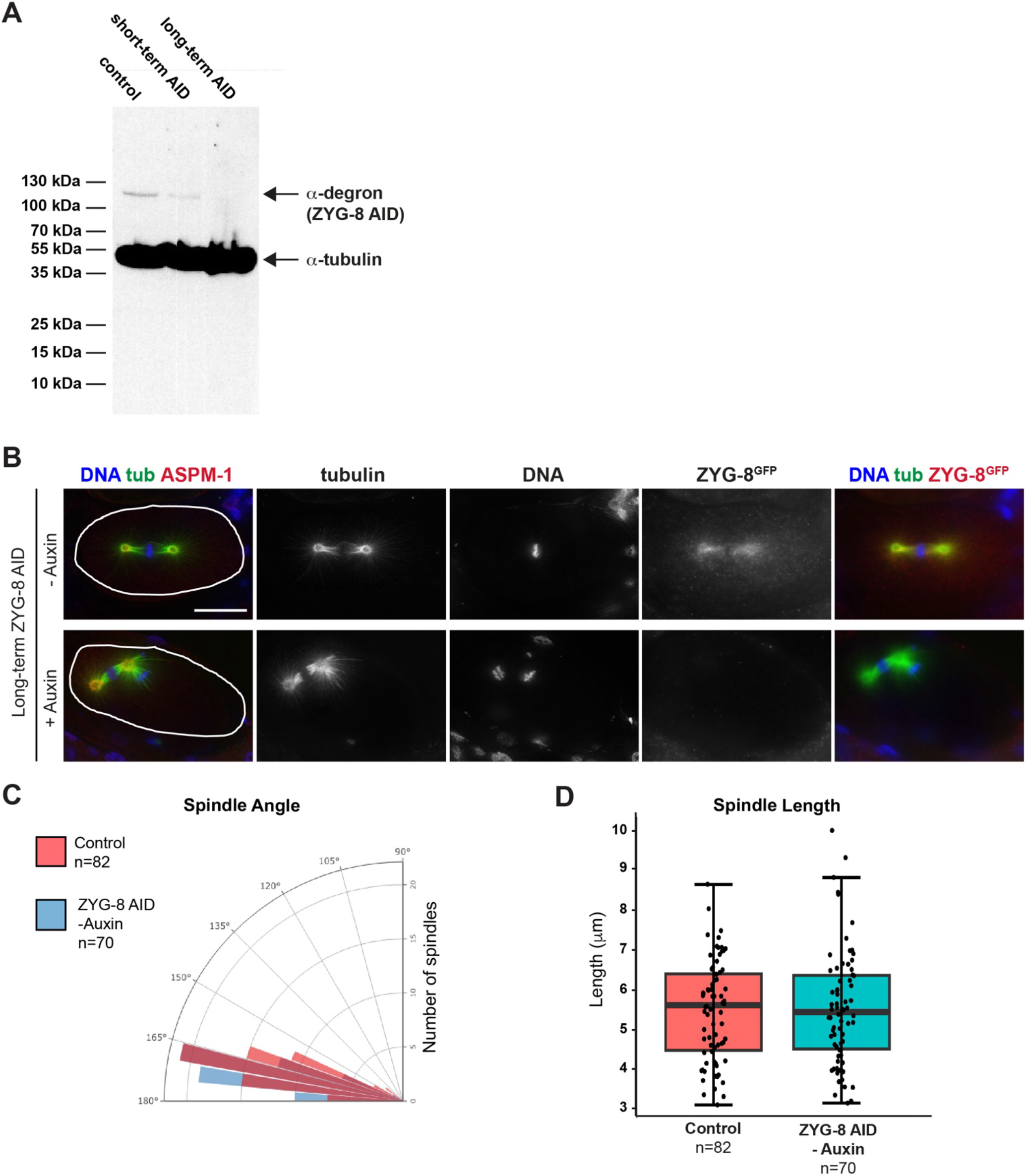
AID allows for temporally and spatially controlled depletion of ZYG-8. (A) Western blot of control, short-term auxin treated (soaking worms for 40 minutes in auxin-containing media), and long-term auxin treated (incubating worms for 18 hours on auxin-containing plates) embryo-only samples. An anti-degron antibody was used to detect ZYG-8 and an anti-tubulin antibody was used as a loading control. (B) IF images of one-cell mitotically dividing embryos in the ZYG-8 AID strain. Auxin treatment resulted in spindle positioning defects (13/15 embryos), phenocopying previous studies of *zyg-8* mutants (Gonczy et al., 2001; Bellanger et al., 2007). (C) Quantification of spindle angle and spindle length in the ZYG-8 AID strain compared to a control strain expressing TIR1 without ZYG-8 tagged; the lengths and angles did not appear significantly different (p>0.5), suggesting that tagging ZYG-8 does not substantially alter protein function. Scale bar = 10µm.

**Figure S3.**
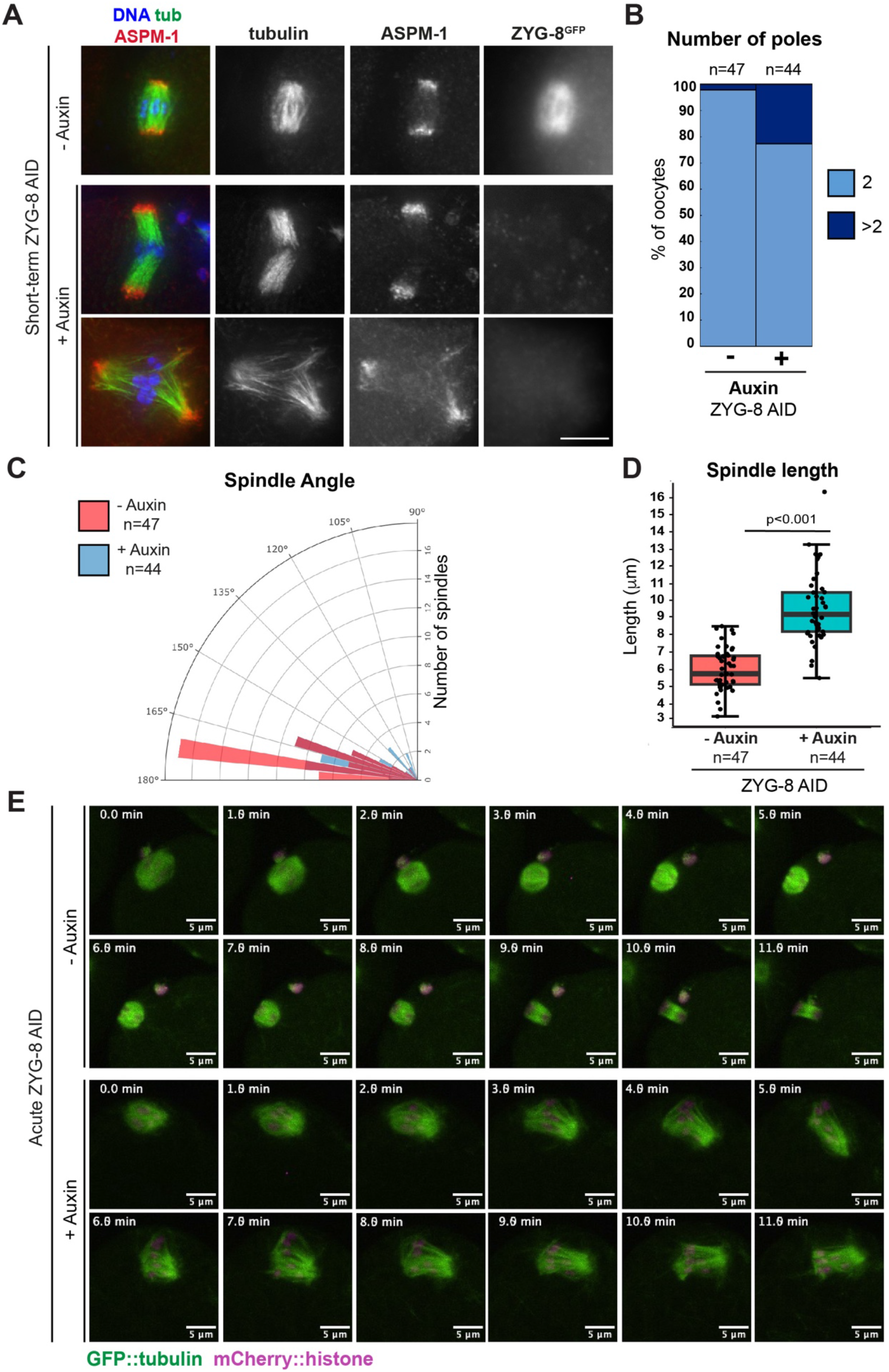
Short-term ZYG-8 depletion in unarrested oocytes reveals the same phenotypes observed following metaphase-arrest. (A) Immunofluorescence images of unarrested oocytes treated with vehicle (row 1) or short-term auxin (rows 2-3); shown are tubulin (green), DNA (blue), and ASPM-1 (red). (B-D) Quantification of the number of ASPM-1-marked poles, spindle angle, and spindle length. Short term ZYG-8 AID results in spindle defects even without metaphase arrest. (E) Live imaging of acute auxin treatment of unarrested spindles; shown are GFP::tubulin (green) and mCherry::histone (magenta). Control spindles maintain bipolarity and eventually segregate chromosomes in anaphase (rows 1-2). In contrast, rows 3-4 show an auxin-treated unarrested spindle elongate and weaken at the midspindle, demonstrating the same defects observed with metaphase arrest. Scale bars = 5µm.

**Figure S4.**
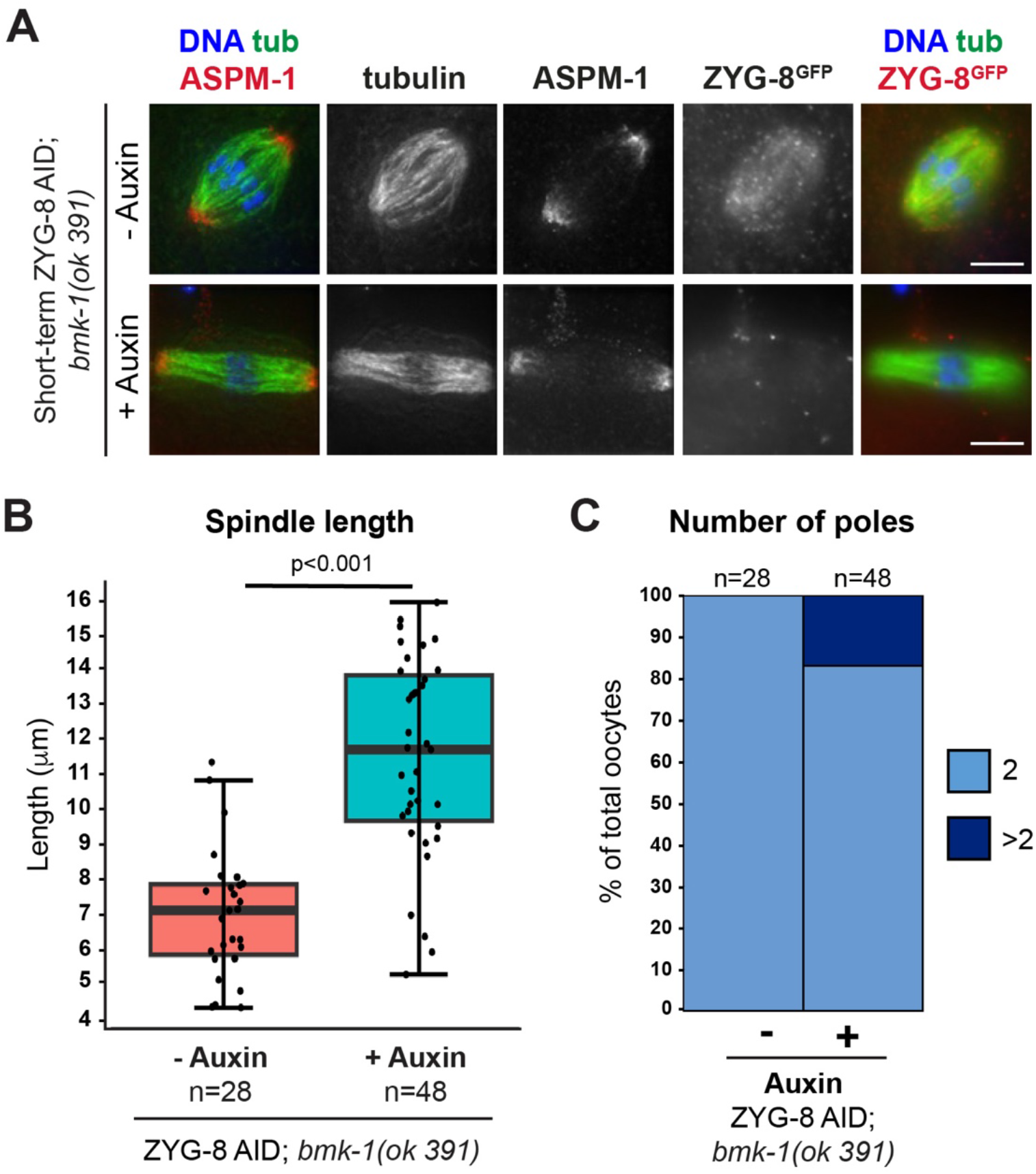
ZYG-8 depletion causes spindle phenotypes in a *bmk-1* mutant without KLP-18 depletion. (A) Immunofluorescence images of oocyte spindles in the ZYG-8 AID; *bmk-1(ok391)* strain in the presence and absence of auxin. Due to diffuse ZYG-8 localization, ZYG-8 images are not deconvolved. (B-C) Quantification of spindle length and the number of ASPM-1-marked poles per spindle. Spindles are longer upon auxin depletion and a fraction of spindles have multiple poles, demonstrating that ZYG-8 likely has functions in addition to regulating BMK-1. Scale bars = 5µm.

## VIDEO LEGENDS

**Video 1. ZYG-8 AID metaphase-arrested oocyte spindles maintain bipolarity in the absence of auxin**

Live imaging of an *emb-30(RNAi)* metaphase-arrested oocyte spindle; corresponds to Figure 3A. Shown are GFP::tubulin (green) and mCherry::histone (magenta). Oocytes were dissected into control Meiosis Medium containing vehicle. The spindle maintains bipolarity; chromosomes oscillate but stay aligned at the midspindle. The phenotype was consistent in all videos (n=5). Scale bar = 5µm.

**Video 2. ZYG-8 AID metaphase-arrested oocyte spindles elongate and lose mid-spindle integrity upon auxin treatment**

Live imaging of an *emb-30(RNAi)* metaphase-arrested oocyte spindle; corresponds to Figure 3A. Shown are GFP::tubulin (green) and mCherry::histone (magenta). Oocytes were dissected into auxin-containing Meiosis Medium. The spindle immediately begins to elongate, the midspindle weakens, and chromosomes lose alignment at the midspindle. The phenotype was consistent in all videos (n=10). Scale bar = 5µm.

**Video 3. ZYG-8 AID unarrested oocyte spindles maintain bipolarity and undergo anaphase in the absence of auxin**

Live imaging of a control oocyte spindle; corresponds to Figure S3E. Shown are GFP::tubulin (green) and mCherry::histone (magenta). Oocytes were dissected into control Meiosis Medium containing vehicle. The spindle maintains bipolarity in metaphase, rotates towards the cortex, shortens, and then elongates as chromosomes segregate bidirectionally. The phenotype was consistent in all videos (n=5). Scale bar = 5µm.

**Video 4. ZYG-8 AID unarrested oocytes exhibit spindle defects following auxin treatment**

Live imaging of an unarrested auxin-treated oocyte spindle; corresponds to Figure S3E. Shown are GFP::tubulin (green) and mCherry::histone (magenta). Oocytes were dissected into auxin-containing Meiosis Medium. The auxin-treated spindle begins to elongate as the midspindle loses integrity, and chromosome become misaligned. Spindle defects were observed in all videos (n=5). Scale bar = 5µm.

**Video 5. *klp-18(RNAi)* spindles maintain a single monopole as chromosomes move towards the center of the aster during anaphase**

Live imaging of a *klp-18(RNAi)* ZYG-8 AID oocyte spindle; corresponds to Figure 5C. Shown are GFP::tubulin (green) and mCherry::histone (magenta). Oocytes were dissected into control Meiosis Medium containing vehicle. Control *klp-18(RNAi)* spindles remain monopolar as chromosomes slowly move towards the center pole in anaphase. The phenotype was consistent in all videos (n=5). Scale bar = 5µm.

**Video 6. Acute ZYG-8 AID causes monopolar spindles to reorganize, reestablish bipolarity, and segregate chromosomes bidirectionally, example 1**

Live imaging of a *klp-18(RNAi)* ZYG-8 AID oocyte spindle; corresponds to Figure 5C. Shown are GFP::tubulin (green) and mCherry::histone (magenta). Oocytes were dissected into auxin-containing Meiosis Medium. Upon treatment with auxin, the monopolar spindle reorganizes into a bipolar spindle that then segregates chromosomes bidirectionally (bidirectional chromosome segregation was observed in 4/12 oocyte spindles). Scale bar = 5µm.

**Video 7. Acute ZYG-8 AID causes monopolar spindles to reorganize and reestablish bipolarity, example 2**

Live imaging of a *klp-18(RNAi)* ZYG-8 AID Meiosis II oocyte spindle; corresponds to Figure 5C. Shown are GFP::tubulin (green) and mCherry::histone (magenta). Oocytes were dissected into auxin-containing Meiosis Medium. Upon treatment with auxin, the monopolar spindle reorganizes and incorporates the polar body, forming a multipolar and finally a bipolar spindle. Monopolar spindles reorganized in 12/12 videos: 3/12 reincorporated the polar body, 6/12 reestablished bipolarity, and 3/12 formed disorganized spindles. Scale bar = 5µm.

**Video 8. ZYG-8 AID; *bmk-1(ok391)* monopolar spindles maintain a single pole and chromosomes move inwards in anaphase in the absence of auxin**

Live imaging of a *klp-18(RNAi)* ZYG-8 AID *bmk-1(ok391)* oocyte spindle; corresponds to Figure 6C. Shown are GFP::tubulin (green) and mCherry::histone (magenta). Oocytes were dissected into control Meiosis Medium containing vehicle. The monopolar spindle maintains a single pole as chromosomes move towards the center of the spindle in anaphase. The phenotype was consistent in all videos (n=5). Scale bar = 5µm.

**Video 9. ZYG-8 AID; *bmk-1(ok391)* monopolar spindles maintain a single pole and chromosomes move inwards in anaphase following auxin treatment**

Live imaging of a *klp-18(RNAi)* ZYG-8 AID *bmk-1(ok391)* oocyte spindle; corresponds to Figure 6C. Shown are GFP::tubulin (green) and mCherry::histone (magenta). Oocytes were dissected into auxin-containing Meiosis Medium. After auxin treatment, the monopolar spindle maintains a single pole as chromosomes move towards the center of the spindle in anaphase. The phenotype was consistent in all videos (n=5). Scale bar = 5µm.

